# The BRUTUS-LIKE E3 ligases BTSL1 and BTSL2 fine-tune transcriptional responses to balance Fe and Zn homeostasis

**DOI:** 10.1101/2022.12.28.522107

**Authors:** Camilla Stanton, Jorge Rodríguez-Celma, Ute Kraemer, Dale Sanders, Janneke Balk

## Abstract

The mineral micronutrients zinc (Zn) and iron (Fe) are essential for plant growth and human nutrition, but interactions between the homeostatic networks of these two elements are not fully understood. Here we show that loss-of-function of *BTSL1* and *BTSL2*, which encode partially redundant E3 ubiquitin ligases that negatively regulate Fe uptake, confers tolerance to Zn excess in *Arabidopsis thaliana*. Double *btsl1 btsl2* mutant seedlings grown on high Zn medium accumulated similar amounts of Zn in roots and shoots as the wild type, but suppressed the accumulation of excess Fe in roots. RNA-seq analysis showed that roots of mutant seedlings had relatively higher expression of genes involved in Fe uptake (*IRT1, FRO2, NAS*) and in Zn storage (*MTP3, ZIF1*). Surprisingly, mutant shoots did not show the transcriptional Fe-deficiency response which is normally induced by Zn excess. Split-root experiments suggested that within roots the BTSL proteins act locally and downstream of systemic Fe deficiency signals. Together, our data show that constitutive low-level induction of the Fe-deficiency response protects *btsl1 btsl2* mutants from Zn toxicity. We propose that BTSL protein function is disadvantageous in situations of external Zn and Fe imbalances, and formulate a general model for Zn-Fe interactions in plants.

**Highlight:** Mutation of two E3 ligases that suppress iron uptake in roots also confers tolerance to zinc toxicity, identifying a regulatory point of interaction between iron and zinc homeostasis.

## INTRODUCTION

Zinc (Zn) and iron (Fe) are two important elements in biology, with plants serving as the primary source of these micronutrients for human nutrition. However, population groups relying upon diets of mainly cereals, which are a poor source of bioavailable Zn and Fe (Cakmak *et al*., 2010), are likely to suffer deficiencies. It is estimated that at least 2 billion people globally are affected, with Fe and Zn deficiency often occurring simultaneously in individuals, leading to a range of health disorders, including anaemia, impaired development and depressed immune system function which can lead to early childhood morbidity (WHO, 2010).

Zn and Fe are found in the *d-*block of the periodic table, alongside other transition metals, such as manganese (Mn) and copper (Cu). Zn and Fe have different chemical characteristics, with Zn found as a stable divalent cation (Zn^2+^), whilst Fe can take on different oxidation states (Fe^2+^ and Fe^3+^) and participate in oxidation-reduction reactions (Habashi, 2013a, b). As such, the ions fulfil very different biochemical roles in enzymes and proteins. However, the divalent cations of Fe^2+^, Zn^2+^ and Mn^2+^ have similar ionic radii and favour similar coordination geometries which can lead to protein mismetallation (Robinson and Glasfeld, 2020). Moreover, these metals can share the same membrane transporters. A particularly well-studied example in plants is the Iron-Regulated Transporter1 (IRT1), the main uptake route of Fe (as Fe^2+^) in the root epidermis, which is also able to transport Zn^2+^, Mn^2+^, Co^2+^, Cd^2+^ and Ni^2+^ (Eide *et al*., 1996; Nishida *et al*., 2011; Vert *et al*., 2002).

The question of how plants coordinate the uptake and distribution of different transition metals in relation to one another has been studied in some detail for Fe and Zn (Hanikenne *et al*., 2021). Physiological studies as well as Arabidopsis mutant studies have provided important insights into the points of cross-talk in the homeostatic networks of these two metals (summarized in **Table 1**), but many questions remain. An early observation is that excess Zn in the soil or medium results in secondary Fe deficiency and Fe deficiency-related phenotypes, such as chlorosis, growth stunting (Fukao *et al*., 2011; Leskova *et al*., 2017; Vert *et al*., 2002) and induction of the transcriptional response to Fe deficiency (Fukao *et al*., 2011; Leskova *et al*., 2017). Increasing the Fe concentration in the medium ameliorates Zn toxicity (Shanmugam *et al*., 2011). These observations could in part be explained by post-translational regulation of IRT1, which helps prevent the uptake of non-Fe metals: a cytosolic loop of IRT1 binds directly to Zn or Mn ions, which leads to internalisation of IRT1 from the plasma membrane and subsequent degradation (Barberon *et al*., 2014; Dubeaux *et al*., 2018). Thus, when Zn levels in the environment are high, excess uptake of Zn into the plant would be prevented by active degradation of IRT1, but this also decreases Fe import. While this mechanism may be effective at moderate Zn levels, at higher concentrations (>50 μM) seedlings do accumulate Zn and display toxicity symptoms.

**Table 1.** Genes and metabolites affecting Zn tolerance in Arabidopsis

| Manipulation | More/less Zn tolerant | Direct effect | Effect on metal distribution |
| --- | --- | --- | --- |
| Increased Fe supply | More | N/A | Increased shoot Fe accumulation and reduced root Zn accumulation (Shanmugam *et al.*, 2011) |
| *fbp* (mutation of *FBP*, *FIT-BINDING PROTEIN*) | More | Increased transcriptional activity of FIT, including de-repression of *NAS1/2/4* in the root stele | Increased retention of Zn in roots and Fe translocation to shoots (Chen *et al.*, 2018) |
| RNAi of *MTP3* (*METAL TOLERANCE PROTEIN 3*) | Less | Reduced expression of the *MTP3*, a vacuole-localized importer of Zn in the roots | Reduced retention of Zn in root vacuoles and increased shoot Zn accumulation (Arrivault *et al.*, 2006) |
| *35S:MTP3* | More | Ectopic expression of the Zn vacuolar importer *MTP3* in rosette leaves | Enhanced Zn accumulation in shoot vacuoles (Arrivault *et al.*, 2006) |
| *HMA3*<sup>Col-0</sup> (*HEAVY METAL ATPASE*, Col-0 accession) | Less | Non-functional variant of vacuolar Zn importer *HMA3* | Reduced Zn retention in root vacuoles (Morel *et al.*, 2009; Park and Ahn, 2017) |
| *nas4-1* (mutation of *NICOTIANAMINE SYNTHASE 4*) | Less | Reduced NA levels in roots and shoots | Lower capacity of Fe uptake and root-to-shoot transport (Palmer *et al.*, 2013) |
| *35S:ZIF1* (*ZINC-INDUCED FACILITATOR 1*) | More | High, ectopic expression of vacuolar NA importer *ZIF1* | Increased NA transport into root vacuoles leading to increased Zn sequestration in roots (Haydon *et al.*, 2012) |
| *zif2* (mutation of *ZINC-INDUCED FACILITATOR 2*) | Less | Reduced expression of vacuolar NA importer *ZIF2* | Decreased NA transport into root vacuoles, leading to reduced Zn sequestration in roots and enhanced translocation to the shoot (Remy *et al.*, 2014) |
| *gsh1* (mutation of *GLUTATHIONE SYNTHASE 1*) | Less | Reduced glutathione biosynthesis | Reduced Fe levels in shoots (Shanmugam *et al.*, 2012) |
| *frd3* (mutation of *FERRIC REDUCTASE DEFECTIVE 3*) | Less | Reduced efflux of citrate into the xylem as a result of loss-of-function of the citrate transporter *FRD3* | 20 μM Zn in the medium rescues root growth and oxidative stress in the *frd3* mutant by competing with Fe in the root cell wall (Scheepers *et al.*, 2020) |
| *FRD3*<sup>Sha</sup> (Shahdara accession) | More | Mild impairment of *FRD3* function leading to reduced efflux of citrate into the xylem | Reduced Fe (and Zn) translocation to shoots (Pineau *et al.*, 2012) |
| *bts-1* (T-DNA insetion in the 5'UTR of *BRUTUS*) | More | Reduced expression of *BTS*, encoding an E3 ubiquitin ligase which targets IVC bHLH transcription factors for degradation | Upregulation of Fe uptake and transport genes, increased Fe accumulation in roots and shoots, increasing tissue Fe/Zn ratio (Zhu *et al.*, 2022) |
| *myb10 myb72* (mutation of Myb transcription factors *MYB10* and *MYB72*) | Less | Reduced expression of *NAS4* and other Fe uptake and transport genes controlled by *MYB10* and *MYB72* | Decreased Fe uptake capacity (Palmer *et al.*, 2013) |

Another important point of cross-talk between Fe and Zn is the <u>F</u>ER-like Iron deficiency induced <u>T</u>ranscription factor FIT (Hanikenne *et al*., 2021). Functioning as a heterodimer together with bHLH38/39/100/101 (group Ib), FIT upregulates genes involved in Fe uptake and translocation such as *IRT1*; genes involved in vacuolar Zn transport; and *Nicotianamine Synthase* (*NAS*) genes required for both processes (Fan *et al*., 2021; Schwarz and Bauer, 2020). Several studies have documented the specific roles of those FIT-regulated genes in Zn homeostasis or Fe-Zn crosstalk, see **Table 1**. In addition, regulators of FIT activity such as FIT Binding Protein (FBP) and the E3 ligase BRUTUS have recently been shown to influence Zn-toxicity phenotypes (summarized in more detail below).

Of the vacuolar Zn transporters regulated by FIT, the *MTP3* gene encoding Metal Tolerance Protein 3 is expressed in root epidermal and cortical cells (Arrivault *et al*., 2006). *MTP3* RNAi lines accumulated higher levels of Zn in shoot tissues under Zn excess and Fe deficiency compared to control lines (Arrivault *et al*., 2006). As such, immobilisation of Zn into root vacuoles by MTP3 decreases Zn mobility in the root and thus limits its root-to-shoot translocation. FIT also upregulates *Heavy Metal ATPase3* (*HMA3*), encoding a vacuolar Zn/Cd/Co/Pb transporter found expressed in the root stele and shoots (Morel *et al*., 2009; Wu *et al*., 2012). The *A. thaliana* Col-0 accession carries a loss-of-function *HMA3* allele, correlated with greater sensitivity to Zn excess due to reduced Zn sequestration in roots (Park and Ahn, 2017).

The FIT-regulated *NAS* genes have a more complex role in Zn and Fe transport and distribution. They are upregulated under Zn excess and Fe deficiency, as well as Zn deficiency (Chen *et al*., 2018; Klatte *et al*., 2009; van de Mortel *et al*., 2006). Nicotianamine (NA) is able to form stable complexes with a range of metals, including Zn^2+^, Fe^2+^ and Mn^2+^ and mediates radial and long-distance metal transport, intracellular transport and vacuolar storage (Bauer and Schuler, 2011; Scholz *et al*., 1992). In addition, T-DNA *nas4* mutants showed reduced NA levels in both roots and shoots and were more sensitive to Fe deficiency, displaying interveinal chlorosis (Koen *et al*., 2013), as well as more sensitivity to excess Zn (Palmer *et al*., 2013). NA additionally plays a role in promoting root Zn sequestration in roots. The Zinc-Induced Facilitator 1 (ZIF1) and 2 (ZIF2) proteins are tonoplast-localised NA transporters that mediate vacuolar localisation of NA-chelated metals in conjunction with MTP3 and HMA3 (Haydon and Cobbett, 2007; Haydon *et al*., 2012; Remy *et al*., 2015; von Wiren *et al*., 1999).

In addition to NA, citrate and glutathione (GSH) are also important chelators shared by Zn and Fe, as well as Mn, for inter- and intracellular transport (Flis *et al*., 2016; Morrissey and Guerinot, 2009; Shanmugam *et al*., 2012). Fe is transported to shoots via the xylem predominantly as Fe^3+^-citrate complexes, and there is also evidence for the presence of Zn-citrate complexes in xylem sap (Cornu *et al*., 2015; Flis *et al*., 2016; Lopez-Millan *et al*., 2000; Rellán-Álvarez *et al*., 2011; White, 1981). Citrate efflux into the xylem is mediated by Ferric Reductase Defective 3 (FRD3), which is induced under Fe deficiency and Zn excess, as well as Zn deficiency (Charlier *et al*., 2015; Durrett *et al*., 2007; Pineau *et al*., 2012). *frd3* mutants showed impaired root-to-shoot Fe transport and constitutive activation of the Fe deficiency response, resulting in very strong accumulation of various secondary substrates of IRT1, in particular Zn and Mn, in shoots (Baxter *et al*., 2008; Delhaize, 1996), as well as accumulation of Fe in cell walls of the vasculature in roots (Durrett *et al*., 2007; Rogers and Guerinot, 2002). Supply of moderate amounts of Zn (20 µM) partially restored root growth by interfering with the accumulation of Fe and hydrogen peroxide in the cell wall (Scheepers *et al*., 2020). The role of GSH in metal homeostasis is still poorly understood, and may be mediated by its direct Fe and metal-binding ability, or its role in scavenging reactive oxygen species in the ascorbate-glutathione cycle, or in nitric oxide signalling upstream of the Fe deficiency response (Shanmugam *et al*., 2015; Xiang *et al*., 2001; Zlobin *et al*., 2017). Loss-of-function *gsh1* mutants, which have reduced GSH levels, show greater sensitivity to Zn excess and are unable to launch an Fe-mediated Zn tolerance phenotype due to diminished root-to-shoot Fe translocation (Shanmugam *et al*., 2012).

While the role of different transporters and chelators in Fe and Zn homeostasis is fairly well characterized, the role of regulatory proteins is only just emerging. The recently discovered FBP inhibits the activity of FIT in the root stele by binding to the DNA-binding domain (Chen *et al*., 2018). *fbp* mutants show increased expression of *NAS* genes and tolerance to Zn excess. FIT protein levels are also regulated post-transcriptionally by the Fe-binding E3 ubiquitin ligases, BRUTUS-LIKE1 (BTSL1) and BTSL2. The *btsl1 btsl2* double mutant fails to downregulate the Fe deficiency response upon Fe resupply, correlating with sustained levels and activity of FIT (Rodriguez-Celma et al., 2019). The closely related BRUTUS (BTS) protein negatively regulates the transcriptional response to Fe deficiency upstream of FIT (Hindt *et al*., 2017; Long *et al*., 2010; Selote *et al*., 2015). Expressed in shoots and the root stele, BTS ubiquitinates the IVc bHLH transcription factors bHLH104, ILR3 and bHLH115 (Kim *et al*., 2019; Long *et al*., 2010; Selote *et al*., 2015), as well as possibly bHLH121 (Kim et al., 2019). RNA-seq analysis of *bts* mutant seedlings showed constitutive upregulation of Fe uptake and transport genes (Zhu *et al*., 2022), enhanced Fe accumulation and tolerance to Fe deficiency (Hindt *et al*., 2017), but it also shows tolerance to Zn excess (Zhu *et al*., 2022). By contrast, the role of BTSL1 and BTSL2 in Zn homeostasis has yet to be explored.

Here we show that *btsl1 btsl2* double mutant seedlings display tolerance to Zn excess (100 µM), partially modulated by the Fe concentration in the medium. RNA-seq analysis, ionomics and split-root experiments suggest that root-specific upregulation of the ferrome, via both FIT-dependent and -independent networks, and moderately increased Fe uptake, mitigates the Zn-induced Fe deficiency response in the mutant. We propose that tight regulation of FIT by the E3 ubiquitin ligases BTSL1 and BTSL2 may have a trade-off in that the homeostatic networks fail when Fe and Zn levels in the soil are very unbalanced.

## Materials and Methods

### Plant materials and growth conditions

*Arabidopsis thaliana* Columbia ecotype (Col-0) plants were used as wild type. T-DNA insertion lines *btsl1-1* (SALK_015054), *btsl2-2* (SALK_048470) and the *btsl1-1 btsl2-2* double mutant (*btsl1x2*) were previously described by (Rodriguez-Celma *et al*., 2019).

Plants were surface sterilised and grown on 0.25 x modified Hoagland medium with 1% (w/v) sucrose and 1.5% (w/v) EDTA-washed agar as described by (Sinclair and Krämer, 2020). The medium was composed of (in mM): Ca(NO_3_)_2_ (1.5), KH_2_PO_4_ (0.28), MgSO_4_ (0.75), KNO_3_ (1.25), CuSO_4_ (0.005), ZnSO_4_ (0.001 or 0.1), MnSO_4_ (0.005), H_3_BO_3_ (0.025), Na_2_MoO_4_ (0.0001) and KCl (0.05). Fe was added in the form of Fe(NO_3_)_3_HBED (N,N′-bis(2-hydroxyphenyl)ethylenediamine-N,N′-diacetic acid), 0.005, 0.05 or 0.1 mM. as indicated. The pH was adjusted to 5.7 with 1 mM 2-(N-morpholino)ethanesulfonic acid (MES). Seedlings were grown vertically on 100 mm^2^ square plates (R & L Slaughter) in a controlled environment room maintained at 23 °C with a photoperiod of 16 h light/8 h dark. For propagation and soil experiments, plants were grown on a peat mix; 600 L Levington F2 peat, 100 L 4 mm grit, 196 g Exemptor® (chloronicotinyl insecticide, GB84080896A, Bayer CropScience, UK) under 22 °C with a photoperiod of 16 h light/8 h dark. Soil was dried and then homogenised with a ZnSO_4_ solution to reach final concentration of 1000 mg/kg DW^-1^ for Zn excess soil growth conditions.

### Growth of plants with split root systems

Seedlings were germinated on 1/4 Murashige and Skoog with 1% (w/v) agar (AGA03, Formedium) and 0.75% (w/v) sucrose buffered to pH 5.7. To generate split roots, at 5-d-old the primary root was excised and seedlings grown for an additional 5 days. Seedlings with two main roots of similar length were transferred to 100 mm^2^ round split plates (Gosselin™, 15438784, Fisher Scientific, UK) containing modified Hoagland medium with either 1 µM ZnSO_4_, 100 µM ZnSO_4_ or 0 µM FeHBED.

### Chlorophyll quantification

Chlorophyll was extracted from dried and powdered 14-d-old seedlings (*ca*. 50 mg). Samples were incubated in 2 mL 100 % (v/v) methanol at room temperature with shaking (350 rpm) for 10 min and then centrifuged for 5 min at 10000 x *g*. 1 mL of solution was transferred to a transparent cuvette on ice and absorbance measured at 650 nm and 665 nm using a spectrophotometer to calculate the total chlorophyll concentration according to (Lichtenthaler, 1987): Chl a + b (μg/ml) = (22.5 x A650) + (4 x A665).

### Inductively Coupled Plasma-Optical Emission Spectrometry (ICP-OES)

For elemental analysis, root and shoot samples were harvested separately (*ca*. 200 mg). Apoplastic cations were desorbed from roots by washing samples in ice cold 2 mM CaSO_4_ and 10 mM EDTA for 10 min followed by 0.3 mM bathophenanthroline disulphonate and 5.7 mM sodium dithionite for 3 minutes as described by (Cailliatte *et al*., 2010). Samples were washed twice with dH_2_0 then dried for 48 h at 65 °C and the dry weight recorded. Dried samples were then digested in 2 mL 65 % (v/v) HNO_3_ and 0.5 mL H_2_0_2_ for 4 h at 95 °C. The digested solution was diluted to 15 mL with ultrapure water and analyzed by ICP-OES (Vista-PRO CCD Simultaneous ICP-OES; Agilent) calibrated with standards: Zn and Fe at 0.2, 0.4, 0.6, 0.8, and 1 mg l^-1^ and Mn at 1, 2, 3, 4, and 5 mg l^-1^. Soft winter wheat flour was used as a reference material (RM 8438; U.S. National Institute of Standards and Technology) and analyzed in parallel with all experimental samples.

### Ferric chelate reductase (FCR) activity

Roots (*ca*. 200 mg) from 14-d-old seedlings were excised, transferred to a 2 mL Eppendorf tube and submerged in 700 µL of assay solution (0.1 mM Fe^3+^-EDTA and 0.3 mM ferrozine). Samples were incubated in the dark for 30 min at room temperature. The solution was then transferred to a cuvette and absorbance at 562 nm was measured using a spectrophotometer (Emre Aksoy and Koiwa, 2013). Root surface FCR activity (μmol Fe^2+^) per gram root fresh weight (gFW) per hour was calculated assuming a molar extinction coefficient of 28.6 mM^-1^ cm^-1^ according to the formula (((A562/28.6)*700)/root FW) * 2.

### Quantitative reverse-transcription PCR (qRT-PCR)

Total RNA was extracted from dried and powdered 100 mg leaf or root tissue sample taken from pooled 14-d-old seedlings using RNeasy Plant Mini Kit (74904, QIAGEN). RNA was then extracted per the manufacturer’s instruction, with a final elution volume of 30 µL, followed by DNase treatment (Turbo DNase kit, Agilent). cDNA synthesis was carried out in 20 µL final volume reactions with 200 − 500 ng mRNA using Invitrogen™ M-MLV reverse transcriptase (28025013, Thermo Fisher Scientific). qRT-PCR was carried out in 20 µL reactions using 20 ng cDNA and SYBR® Green JumpStart™ ReadyMix™ (S4438, Sigma-Aldrich). *ACTIN2* and *TIP41* were used as reference genes and relative gene expression performed using 2^−ΔΔCt^ method (Livak and Schmittgen, 2001).

### RNA-sequencing (RNA-seq)

RNA was extracted from separated shoots and roots of 14-d-old seedlings using RNeasy Plant Mini Kit (74904, QIAGEN). Quality check was carried out on a 2100 Bioanalyzer Instrument (G2939BA, Agilent Technologies) using the Agilent RNA 6000 Nano Kit (5067-1511, Agilent Technologies). Samples were diluted to 100 ng/µL and a total volume of 30 µL was sent for sequencing. Construction of cDNA sequencing libraries and next generation sequencing (NGS) was carried out by Novogene (Hong Kong). Libraries were generated using NEBNext® Ultra™ RNA Library Prep Kit for Illumina (NEB) and then sequencing run on an Illumina Novaseq 6000 (Illumina, Ca, USA) using 150 bp paired end reads. Quality control was carried out on raw reads using FastQC (Andrew, 2010).

### Read alignment and differential gene expression (DGE)

Reads were pseudoaligned to the TAIR10 transcriptome (TAIR10_cdna_20101214_updated.fa) and transcript abundance calculated using Kallisto (v 0.46.1) (Bray *et al*., 2016) on the NBI high-performance computing (HPC) platform (NBI, Norwich, UK). DGE analysis was then performed using Sleuth (Pimentel *et al*., 2017) in R Studio with a 2-fold (log_2_FC > 1) cut off and a 5 % false discovery rate (adjusted p-val, qval < 0.05) using pairwise comparisons between samples. Sleuth was also used to run principal component analysis. Kallisto and Sleuth were both run in R Studio. Results were verified by performing DGE analysis with EdgeR on the Degust platform (Powell, 2015) and the top results compared.

### Bioinformatic analysis of RNA-seq data

Shoot and root data were analysed separately. Gene list comparisons and Venn diagrams were generated using Gene List Comparator (Cermak, 2020). Hierarchical clustering of differentially expressed genes based on FC difference from the mean was performed using Morpheus (https://software.broadinstitute.org/morpheus) based on Pearson correlation and average linkage method, as well as generation of Heatmaps. Gene Ontology (Mi *et al*., 2019) for biological processes and KEGG pathway (Kanehisa, 2019; Kanehisa and Goto, 2000; Kanehisa *et al*., 2019) enrichment analysis was performed using David (Database for Annotation, Visualization and Integrated Discover) (Huang da *et al*., 2009a, b) filtered with an adjusted p-value (Benjamini < 0.05).

## RESULTS

### *btsl1 btsl2* double mutant seedlings are tolerant to excess Zn

Previous work showed that the *btsl1 btsl2* double knockout mutant (further called *btsl1x2*) is more tolerant to Fe deficiency and more sensitive to Fe excess than wild-type seedlings, correlating with constitutively induced Fe uptake in the mutant (Hindt *et al*., 2017; Rodriguez-Celma *et al*., 2019). *btsl1x2* mutants also accumulated more Zn but not more Mn (Hindt *et al*., 2017), however growth in response to these divalent metal ions has not been investigated.

First, we compared growth phenotypes of the *btsl* single mutants and the *btsl1x2* double mutant on medium with a standard level of Zn (1 μM ZnSO_4_) or with Zn excess (100 μM ZnSO_4_), containing the same concentration of Fe (5 µM FeHBED). The toxic effect of Zn on Arabidopsis seedlings manifested itself in chlorosis and decreased biomass (**Figure 1**) as previously shown (Fukao *et al*., 2011; Leskova *et al*., 2017; Vert *et al*., 2002). In 14-day-old wild-type seedlings grown on excess Zn, the levels of chlorophyll were 30% compared to seedlings grown on standard medium (**Figure 1B**). By contrast, the *btsl1x2* mutant retained 2-fold more chlorophyll than wild type in the presence of 100 µM Zn (570 ± 57 vs 300 ± 13 μg chlorophyll g^-1^ FW, p < 0.01, **Figure 1B**). Both the *BTSL1* and *BTSL2* genes contribute to Zn-induced chlorosis, as no significant increase in chlorophyll was observed in the single mutants (**Supplementary Figure S1B**). Increasing the Fe concentration to 50 or 100 µM, known to suppress the effects of Zn toxicity in wild-type seedlings (Shanmugam *et al*., 2011), further restored chlorophyll levels in *btsl1x2* (**Figure 1B**).

**Figure 1:**
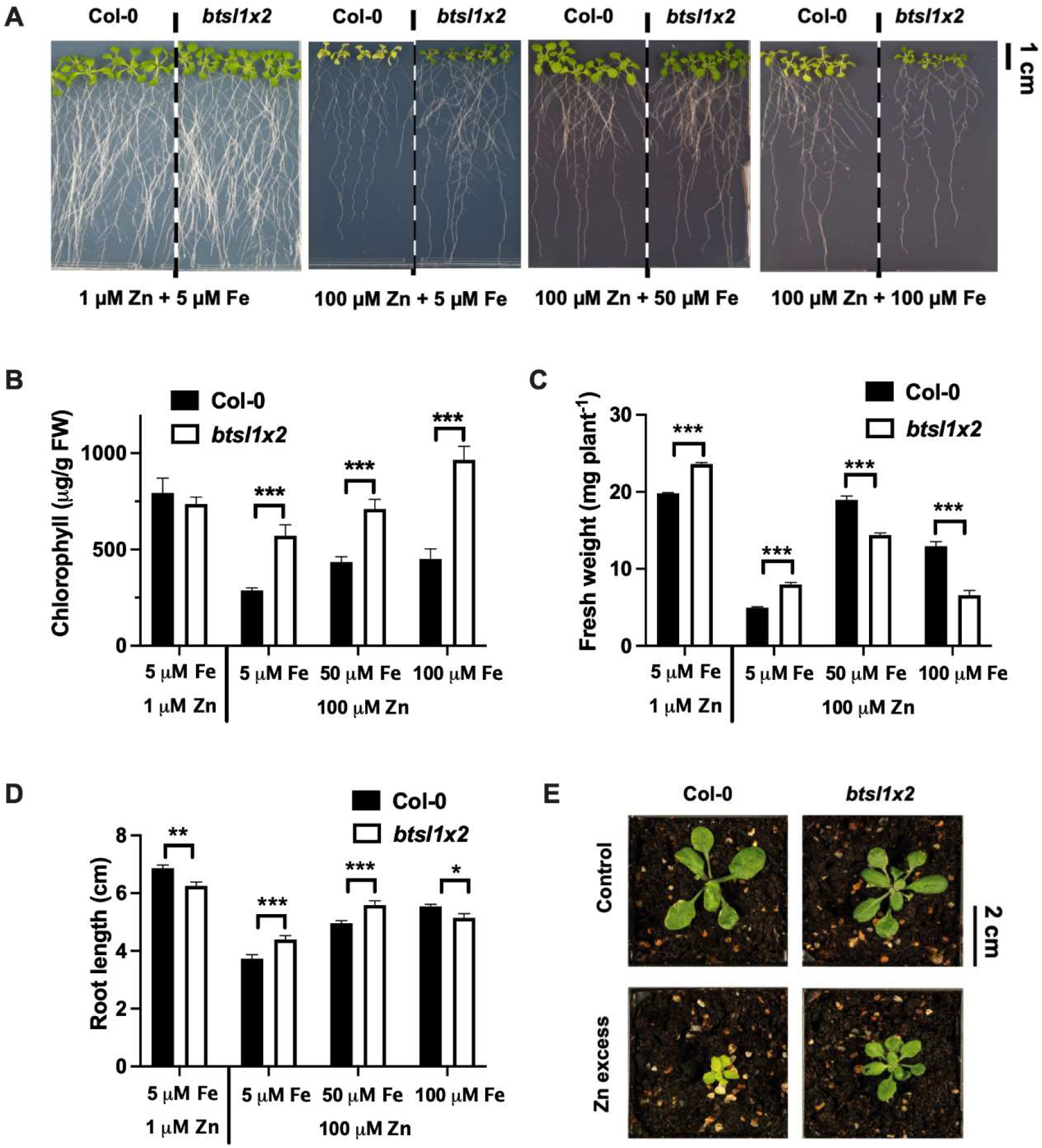
The *btsl1 btsl2* double mutant displays Zn tolerance, partially modulated by Fe in the medium. A) Representative photographs of 14-day-old seedlings of wild type (Col-0) and *btsl1 btsl2* double mutant (*btsl1x2*), grown on modified Hoagland medium with 1 μM ZnSO_4_ (standard conditions) or 100 μM ZnSO_4_ (excess), in the presence of 5, 50 or 100 μM FeHBED as indicated. B) Total chlorophyll normalized to fresh weight (FW) in shoots from seedlings in (A). C) Shoot biomass, in mg fresh weight (FW), of seedlings in (A). D) Primary root length of 10-day-old seedlings. E) Representative photographs of Col-0 and *btsl1x2* plants grown for 22 days on soil without (Control) and with added ZnSO4 (Zn excess) at a concentration of 1 g per kg dry weight). For B – D, data represent mean values (± SEM) from three independent experiments, each comprising six plants per genotype and condition. Statistically significant differences are indicated by asterisks (*p < 0.05, **p < 0.01, ***p < 0.001) as determined by two-way ANOVA followed by Tukey HSD post-hoc test. Non-significant differences are not indicated.

Shoot biomass was higher in *btsl1x2* than in wild type exposed to 1 or 100 µM Zn (120% and 160%, respectively), but lower than wild type at higher Fe concentrations (**Figure 1C**). Root growth was inhibited in the *btsl1x2* mutant on standard medium, most likely due to toxic effects of Fe accumulation (Hindt *et al*., 2017; Rodriguez-Celma *et al*., 2019). Under Zn excess, *btsl1x2* seedlings grew longer roots than wild type in the presence of 5 and 50 µM Fe, but not with 100 µM Fe (**Figure 1D**). The higher chlorophyll concentration and shoot biomass were also evident in the *btsl1x2* mutant when grown on soil with excess Zn (1000 mg ZnSO_4_ per kg soil) (**Figure 1E**).

Zn and Fe homeostasis has also been shown to be interlinked with the homeostasis of other divalent cations, such as Mn^2+^ (Rai *et al*., 2021). To test the growth response of the *btsl1x2* mutant to Mn, seedlings were grown on increasing MnSO_4_ concentrations ranging from Mn deficiency (0 μM MnSO_4_) to excess (250 μM MnSO_4_ (**Supplementary Figure S2A**). There were no significant differences in chlorophyll levels between genotypes or Mn concentrations (**Supplementary Figure S2B**). The *btsl1x2* mutant had more shoot biomass on 0 μM Mn but less biomass on 50 and 250 μM Mn compared to wild type (**Supplementary Figure S2C**), and shorter root lengths on 5 and 50 μM Mn (**Supplementary Figure S2D**). Thus, the *btsl1x2* mutant was more tolerant to Mn deficiency but less tolerant to excess, similar to its growth response to Fe.

Together, these data show that loss of both *BTSL1* and *BTSL2* gene expression results in enhanced tolerance to Zn excess, with further suppression of toxicity effects when higher concentrations of Fe are supplied to the medium, until this also impairs growth.

### *btsl1x2* seedlings under Zn toxicity accumulate similar amounts of Zn but much less Fe in roots than wild type

To explore whether the BTSL proteins affect the distribution of Zn, Fe or Mn between roots and shoots, elemental analysis was carried out by ICP-OES on 14-day-old seedlings grown on medium with standard or excess Zn. Roots were washed with EDTA to ensure desorption of apoplastic cations. There was no significant difference in the shoot or root Zn concentrations between wild type and *btsl1x2* under standard growth conditions (**Figure 2A**). As expected, the Zn concentration in both shoots and roots increased dramatically in wild-type seedlings exposed to 100 µM ZnSO_4_ (23-fold and 89-fold respectively) and by a similar amount in the *btsl1x2* mutant (17-fold and 69-fold respectively).

**Figure 2:**
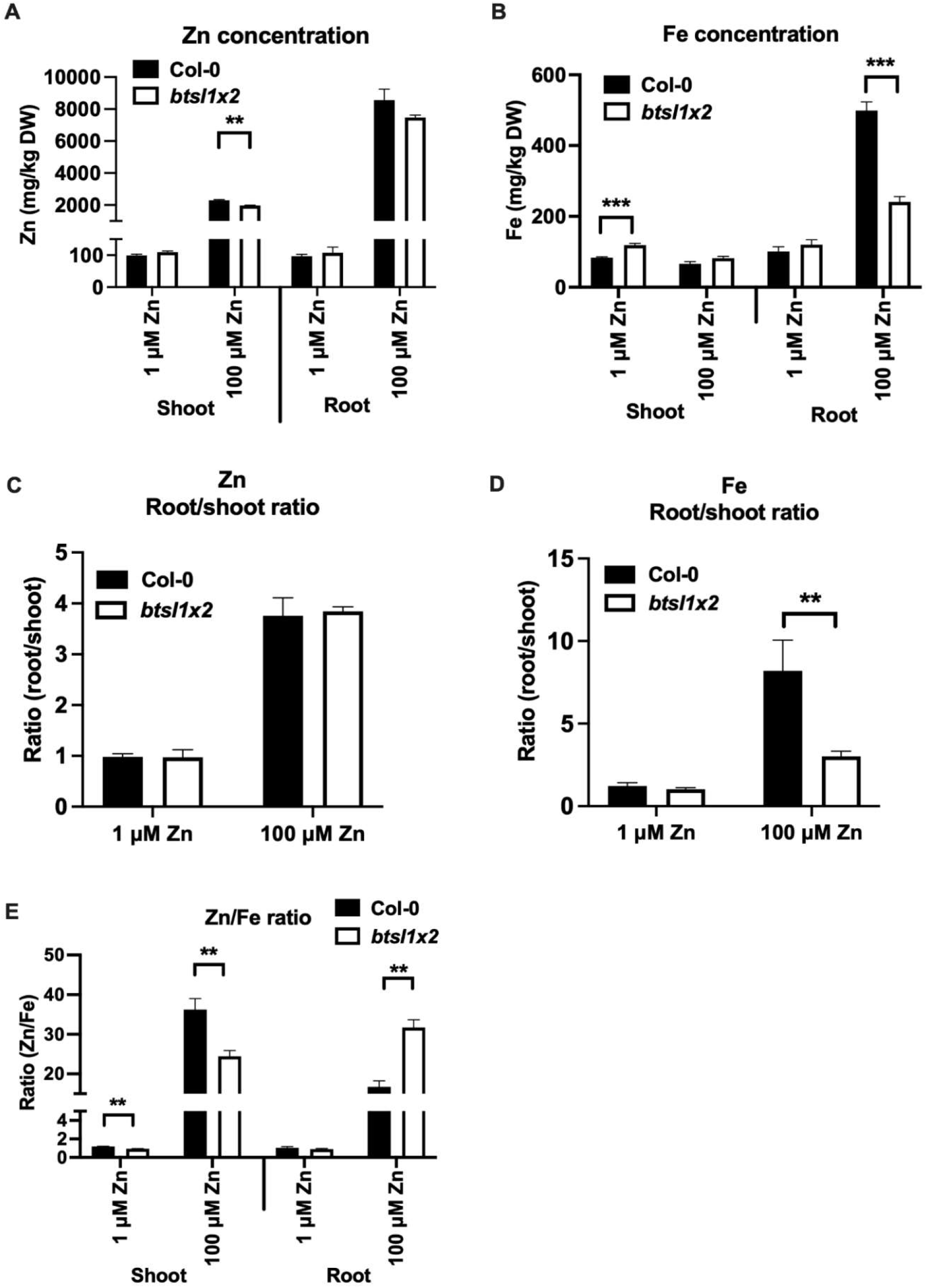
Distribution of Fe and Zn between roots and shoots in response to excess Zn. The concentration of A) Zn and B) Fe in the roots and shoots of 14-d-old seedlings of wild type (Col-0) and *btsl1 btsl2* double mutant (*btsl1x2*) grown on standard (1 μM) or Zn excess (100 μM). Roots were desorbed with chelators to remove apoplastic cations prior to element analysis. Data represent mean values (± SEM) from three independent experiments, each comprising ten plants per genotype and condition. Statistically significant differences are indicated by asterisks (*p < 0.05, **p < 0.01, ***p < 0.001) as determined by two-way ANOVA followed by Tukey HSD post-hoc test. Non-significant differences are not indicated. C) Root-to-shoot ratio of Zn and D) Fe. E) Zn-to-Fe ratio of shoots and roots.

Consistent with previous reports (Hindt *et al*., 2017; Rodriguez-Celma *et al*., 2019), *btsl1x2* seedlings accumulated 1.4-fold more Fe in shoots compared to wild type when grown under standard conditions (120 ± 5 vs 80 ± 2 mg Fe kg^-1^, p < 0.001), whereas the roots contained similar Fe concentrations as wild type (**Figure 2B**). Induction of the Fe deficiency response by excess Zn led to 5-fold Fe accumulation in wild-type roots, and very little of this Fe was translocated to shoots. The *btsl1x2* mutant also accumulated Fe in the roots when grown under Zn excess, but only about half as much as wild-type roots (240 ± 15 vs 500 ± 24 mg Fe kg^-1^, p < 0.01, **Figure 2B**). The calculated root/shoot ratio of Zn was similar in wild type and *btsl1x2*, however the Fe root/shoot ratio was 8 in wild type and 3 in *btsl1x2* (**Figure 2C and D**). A reduced accumulation of Fe in roots and lower Fe root/shoot ratio was previously observed in the *fbp* mutant under Zn excess (Chen *et al*., 2018). Conversely, the *bts* mutant showed increased root Fe accumulation under Zn excess compared to wild type (Zhu *et al*., 2022). In terms of the Zn/Fe ratio, this was lower in the shoots of *btsl1x2* compared to wild-type seedlings (25 vs 37, **Figure 2E**) and, together with the higher chlorophyll levels and shoot biomass in the mutant, indicates that the relatively small trend towards a more physiological Zn/Fe balance may suppress the Zn toxicity effects when Fe is the limiting nutrient (**Figure 1**). By contrast, the root Zn/Fe ratio was relatively higher in the *btsl1x2* mutant while root length was increased compared to wild type.

Under standard growth conditions, there was no significant difference in Mn shoot or root accumulation between the *btsl1x2* mutant and wild type (**Supplementary Figure S2E**). Under Zn excess, the Mn concentrations were reduced in both genotypes, particularly in the roots, with only minor differences between the genotypes. The Mn root/shoot ratio was reduced in both genotypes under Zn excess, but there was no difference between genotypes (**Supplementary Figure S2F**).

In summary, the most striking difference in metal distribution in high Zn conditions is that *btsl1x2* seedlings exhibit suppressed Fe accumulation in the roots, with only minor effects on Zn, Fe or Mn concentrations in the shoots.

### Distinct transcriptomic changes are observed in *btsl1x2* roots, but not shoots, in response to Zn excess

To investigate changes in gene expression that could underpin Zn tolerance in the *btsl1x2* mutant, RNA-seq analysis was carried out separately on roots and shoots of seedlings grown on standard (1 μM ZnSO_4_) and high Zn (100 μM ZnSO_4_) medium (**Figure 3; Supplementary Figure S3**). Principal component analysis (PCA) of the RNA-seq data showed good reproducibility of the data for the three biological replicates of each genotype and treatment (**Figure 3A; Supplementary Figure S3A**). The gene expression in wild-type and *btsl1x2* roots clustered closely together under standard conditions but diverged when exposed to excess Zn (**Figure 3A**).

**Figure 3:**
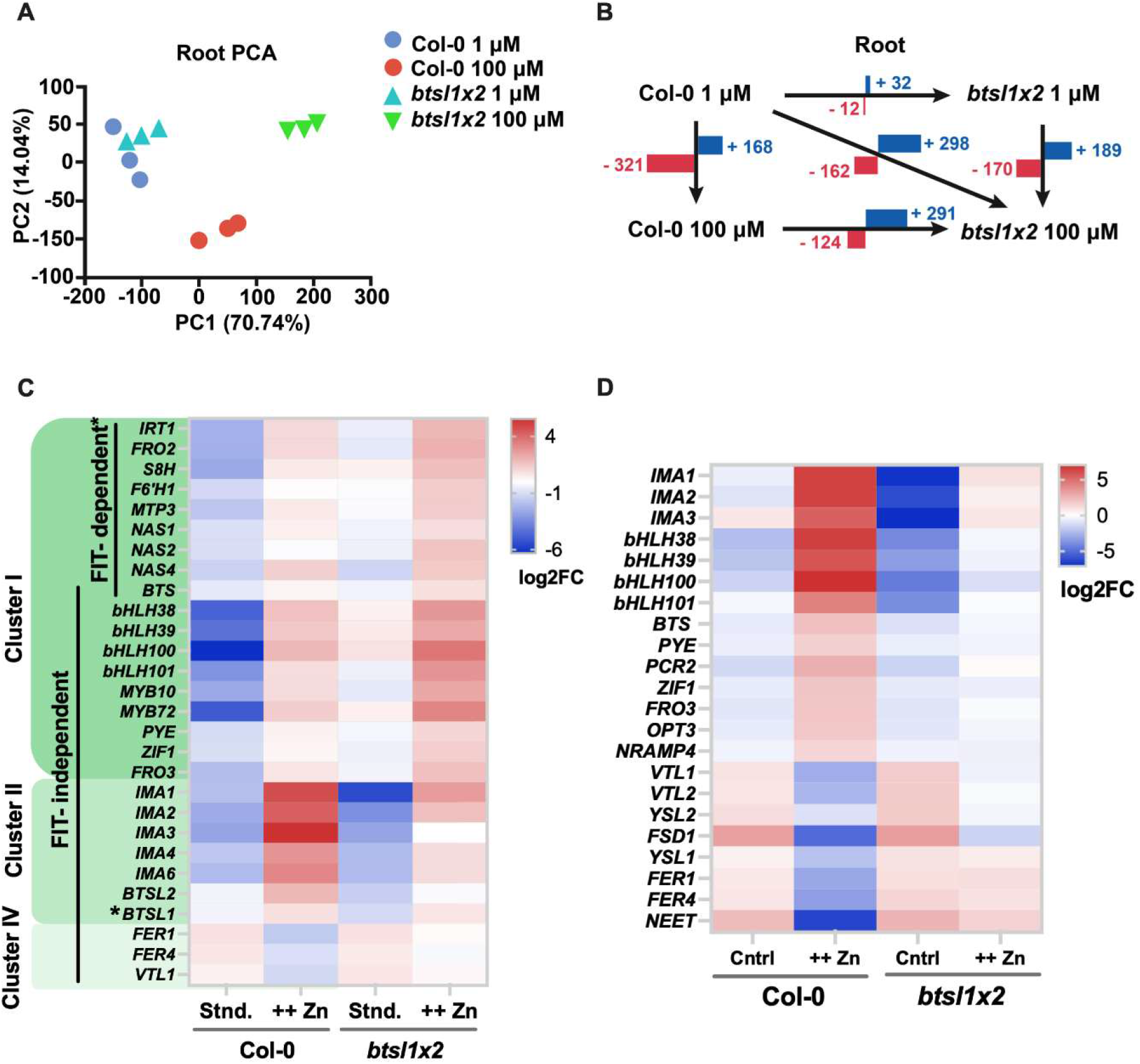
The *btsl1 btsl2* double mutant has a distinct root transcriptomic response to Zn excess. RNA-seq analysis of root samples of wild type (Col-0) and *btsl1 btsl2* (*btsl1x2*) double mutant seedlings, grown for 14 days on standard medium or with excess Zn (100 µM ZnSO_4_). A) Principal component analysis (PCA) of root sample transcripts PC1 = principle component 1, PC2 = principle component 2, (%) is the percentage of variance explained by each PC. B) Number of significantly (adjusted p-value > 0.05, log2(fold change) ≥ 1) upregulated (+) or downregulated (-) genes between all pairwise comparisons. C) Heatmap of selected genes, from a core set of Fe deficiency-responsive genes as defined in Mai et al., 2016, in roots and D) shoots. Expression is represented as standard deviation from the mean global gene expression of pooled replicates. Hierarchical clustering of all differentially expressed genes is shown in Suppl Figure 5 and Suppl Figure 8.

Consistent with close clustering of samples, compared with wild type, only 44 genes were differentially expressed in *btsl1x2* roots under standard conditions (32 upregulated, 12 downregulated, **Figure 3B**). Of this gene set, 28 of those upregulated (including *bHLH38, bHLH39, bHLH100, bHLH101, MYB10, MYB72* and *FRO3*) and 9 of those downregulated (including abiotic stress-responsive *CYB7B122* and flowering-time regulator *FLC*) were also differentially expressed in response to Zn excess, suggesting that a number of Zn excess-associated genes are constitutively more highly expressed in *btsl1x2*.

Comparing the mutant and wild type under Zn excess, 460 genes were differentially expressed in roots (298 upregulated, 162 downregulated, **Figure 3B**) and 771 genes in shoots (207 upregulated and 564 downregulated, **Supplementary Figure S3B**). Under Zn excess compared with standard conditions, 489 genes were differentially expressed in wild-type roots, and 359 in *btsl1x2* roots (**Figure 3B**). By contrast, only 29 genes were differentially expressed in the shoots of the *btsl1x2* mutant in response to excess Zn, whereas wild-type shoots had 1277 differentially expressed genes (**Supplementary Figure S3B; Figure 3D**). This suggests that mutation of the *BTSL1* and *BTSL2* genes, which are expressed in the roots, shields the shoots from the toxic effects of excess Zn in the medium.

### *btsl1x2* roots have induced expression of Fe homeostasis and Zn storage genes under standard conditions and excess Zn, but lower expression of *IMA* genes

Gene Ontology (GO) enrichment analysis showed that in roots, genes associated with Zn and Fe homeostasis were significantly upregulated in both genotypes under Zn excess, whilst commonly downregulated genes were enriched for terms associated with oxidative and biotic stress (**Supplementary Figure S4A**). Genes that were uniquely upregulated in the *btsl1x2* mutant under both standard conditions and Zn excess are associated with Fe homeostasis and transcriptional regulation, in agreement with the demonstrated role of BTSL2 in regulating activity of the FIT transcription factor (Rodriguez-Celma *et al*., 2019).

To further explore how gene expression was impacted in the *btsl1x2* mutant, hierarchical clustering of all the significant DEGs in roots was performed (**Supplementary Figure S5**) and the expression pattern of a well-documented core set of genes involved in Fe and Zn homeostasis (Gao *et al*., 2019; Mai *et al*., 2016; van de Mortel *et al*., 2006) was investigated (**Figure 3C**). Cluster I contained genes that were strongly upregulated in both genotypes under Zn excess but were expressed more highly in the mutant than wild type under both standard and Zn excess conditions. FIT-regulated genes, such as *IRT1, FRO2, MTP3* and *NAS1/2/4* were found in this cluster, consistent with the accumulation of FIT protein in the *btsl1x2* mutant (Rodriguez-Celma *et al*., 2019). Interestingly, FIT-independent genes were also found in this cluster, including FIT-interacting partners *Ib bHLH* genes (*bHLH38/29/100/101*), *MYB10/72, PYE, FRO3* and the Zn tolerance-associated *ZIF1* (**Figure 3C**), in support of the suggestion that other bHLH transcription factors may be targeted by BTSL1 and BTSL2 (Lichtblau *et al*., 2022). Increased expression of Fe homeostasis genes, including *IRT1, Ib bHLH* genes and *FRO2*, has also been observed in the *bts-1* mutant under Zn excess compared to wild type, although the genes were not upregulated under standard conditions (Zhu *et al*., 2022). Moreover, increased expression of *NAS* genes was previously shown to underlie the Zn tolerance phenotype of the *fbp* mutant, although other Fe deficiency-responsive genes were downregulated in *fbp* (Chen *et al*., 2018) that are not down in *btsl1x2*.

Cluster II contained genes that showed generally lower levels of expression in the *btsl1x2* mutant under standard conditions, and were also upregulated in response to Zn excess in both genotypes, but to a lesser degree in the mutant than in the wild type (**Figure 3C**). Fe deficiency-induced *IMA* signalling peptides were found in this cluster. The phloem-mobile *IMA* signalling peptides are thought to induce Fe deficiency transcriptional responses in the roots by inhibiting BTS and BTSL protein interactions with IVb and IVc bHLH transcription factors (Li *et al*., 2021; Lichtblau *et al*., 2022). Reduced expression of *IMA* genes in *btsl1x2* is consistent with the reduced Zn toxicity symptoms and transcriptomic response in shoots (**Supplementary Figure S4 and S6**). As expected, the expression of *BTSL1* and *BTSL2* was also reduced in the mutant, serving as an internal control for the data set. Cluster IV contained genes that were downregulated in both genotypes in response to Zn excess but were more Zn-responsive in wild type than the mutant (**Figure 3C**). Fe storage ferritin genes *FER1* and *FER4*, and the vacuolar Fe transporter, *VTL1*, were found in this cluster.

To verify some of the RNA-seq data, qRT-PCR analysis was performed of selected genes *bHLH38* and *bHLH39*, **(Supplementary Figure S7A – B)**, *ZIF1* and *FRO3* (**Supplementary Figure S7C – D**) and *IRT1* and *FRO2* (**Supplementary Figure S7E – F**) in root samples. The expression of the six genes was enhanced in the *btsl1x2* mutant under Zn excess, in agreement with RNA-seq data.

In shoots, genes uniquely upregulated in wild type in response to Zn were associated with oxidative and biotic stress, whilst downregulated genes were involved in photosynthesis (**Figure 3D** and **Supplementary Figure S6)** in agreement with growth impairment and chlorosis (**Figure 1**). Furthermore, major Fe homeostasis genes, such as *Ib bHLH* genes, *BTS* and *PYE* were strongly upregulated in wild type, but not in the mutant (**Figure 3D** and **Supplementary Figure S8**). This confirms that wild-type shoots respond to Zn as if they are physiologically Fe-deficient, but surprisingly *btsl1x2* shoots do not have this response. However, *IMA* transcripts encoding signalling peptides were partially upregulated in mutant shoots, and *VTL1/2* transporters partially downregulated, in response to Zn (**Figure 3D and Supplementary Figure S8**). While this could indicate that some iron deficiency is sensed in *btsl1x2* shoots, for example in a specific tissue or cell type, it is also possible that the BTSL-FIT module affects these genes indirectly.

Together, transcriptomic analysis shows that BTSL1 and BTSL2 negatively regulate the expression of Fe homeostasis and Zn tolerance-associated genes of both FIT-dependent and -independent networks. Under Zn excess, the transcript levels of genes for Fe uptake, NA biosynthesis and Zn storage are induced in the *btsl1x2* mutant roots, but not in shoots, except for *IMA* paralogs.

### Ferric chelate reductase (FCR) activity is suppressed under Zn excess in wild-type roots compared to *btsl1x2* roots

Whilst increased expression of Fe uptake genes in the *btsl1x2* mutant is consistent with increased Fe accumulation in the shoots under standard growth conditions, it does not explain why Fe accumulation in the roots is suppressed under Zn excess (**Figure 2B**). To see if Fe uptake capacity in the mutant was impacted post-transcriptionally under Zn excess, ferric chelate reductase (FCR) activity was measured and compared to *FRO2* transcript levels. In wild-type roots, FCR activity is induced under both Fe deficiency and Zn excess, but the magnitude of induction is 2-fold lower under Zn excess compared to Fe deficiency (**Figure 4A**) despite comparable transcript levels (**Figure 4B**). In the *btsl1x2* mutant, FCR activity is similar in roots grown under Fe deficiency or Zn excess, and thus 2-fold higher than in Zn-exposed wild-type roots. This finding suggests that in wild-type seedlings, FCR activity mediated by FRO2 is suppressed post-transcriptionally due to excess Zn, which might lead to the observed accumulation of a pool of Fe that is not removed by washing with EDTA (**Figure 2**).

**Figure 4:**
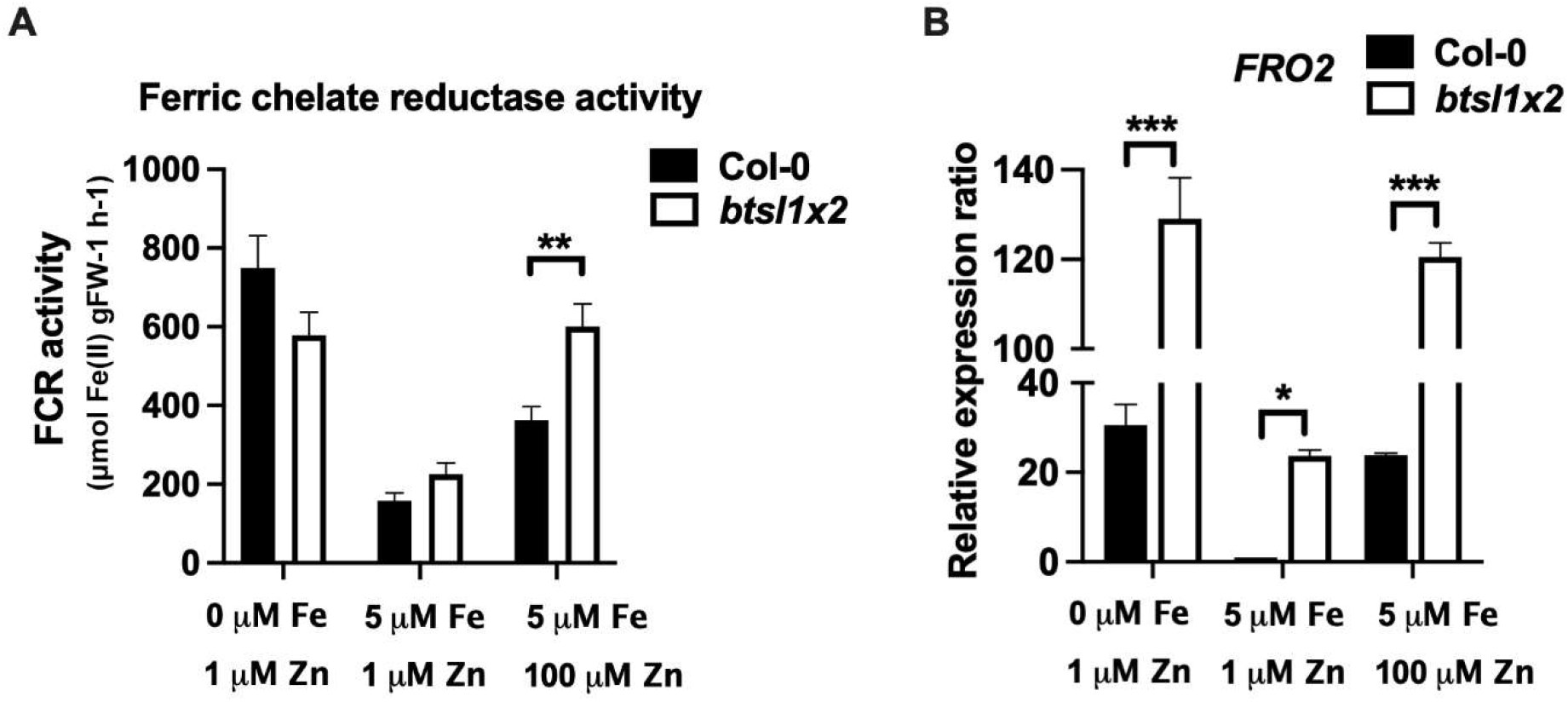
The *btsl1 btsl2* double mutant has higher FCR activity than wild type under Zn excess. A) Ferric chelate reductase (FCR) activity and (B) transcript levels of *FRO2*, the gene responsible for FCR activity on the root surface under Fe deficiency. Wild type (Col-0) and *btsl1 btsl2* mutant (*btsl1x2*) seedlings were grown on medium lacking Fe (0 μM FeHBED, 1 μM ZnSO_4_), standard conditions (5 μM FeHBED, 1 μM ZnSO_4_) or excess Zn (5 μM FeHBED, 100 μM ZnSO^4^) for 10 days, after which roots were excised for the activity assays and qRT-PCR. Fold change is relative to Col-0 and normalised to reference genes *ACTIN2* and *TIP41*. Data represent mean values (± SEM) from three independent experiments, using 5 seedlings per measurement.. Statistically significant differences are indicated by asterisks (*p < 0.05, **p < 0.01, ***p < 0.001) as determined by two-way ANOVA followed by Tukey HSD post-hoc test.

### BTSL proteins act locally in Zn-exposed or Fe-deficient roots

To investigate the contribution of both local and systemic signals to the Zn-induced transcriptional response, we performed split-root experiments (**Figure 5A**). Similar experiments were instrumental in demonstrating that apoplastic Fe causes local induction of Fe-uptake genes as well as a shoot-to-root signal acting systemically on the root transcriptional response (Vert *et al*., 2003), later confirmed by many other studies. As a marker of Zn-induced changes in both shoots and roots, the differentially expressed, FIT-independent, *bHLH38* gene was selected based on our RNA-seq analysis (**Figure 3C**) and quantified by RT-qPCR. First, we verified the effect of exposing half of the root system to Fe deficiency on *bHLH38* expression in the shoots, as this has not been reported in the literature to our knowledge, but the information would be useful for comparison to the Zn-excess response. Seedlings were germinated on standard medium for 10 days, then transferred to split agar plates, with one half of the root system exposed to standard conditions (5 μM FeHBED) and the other half to Fe deficiency (0 μM) for another 7 days (**Figure 5B**). This situation resulted in only a slight increase in *bHLH38* transcript levels in the shoots in both wild type and *btsl1x2* (6-fold and 2-fold respectively), compared to a dramatic 300-fold increase in *bHLH38* expression when the whole root system of wild-type seedlings is Fe deficient. Thus, the availability of Fe to one half of the root system prevents a transcriptional Fe-deficiency response in the shoots.

**Figure 5:**
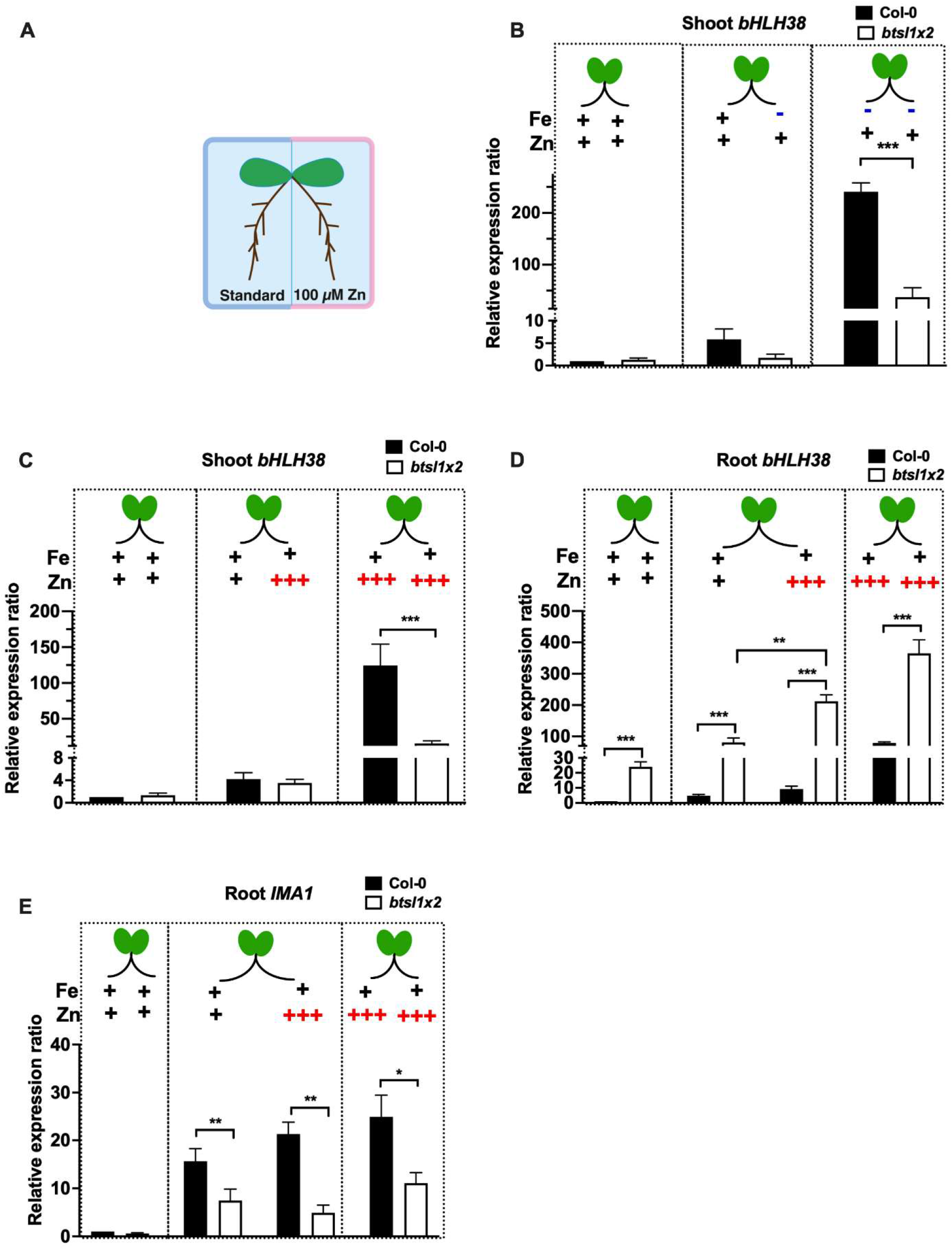
Systemic Fe signalling in *btsl1 btsl2* mutant seedlings in response to Fe deficiency and Zn excess. Wild type (Col-0) and *btsl1 btsl2* mutant (*btsl1x2*) seedlings were germinated on standard medium (1 μM ZnSO4). The primary root was excised at day 5 to promote development of lateral roots. At 10 days, seedlings were transferred to split agar plates with the indicated Fe and Zn concentrations. Samples for RNA extraction were taken 7 days later. A) Schematic showing the experimental set up of the split-root assay. B, C) Expression of *bHLH38* in shoots in response to the roots being exposed to (B) Fe deficiency or (C) Zn excess. D, E) Expression of (D) *bHLH38* and (E) *IMA1* in roots in response to Zn excess exposure in different halves of the roots system. Fold change is relative to Col-0 control and normalised to reference genes *ACTIN2* and *TIP41*. Statistical analysis as in Figure 4B.

When half the root system was exposed to Zn (**Figure 5C**), the expression of *bHLH38* was similarly not highly induced in the shoots, nor in Zn-exposed roots of wild type. By contrast, *bHLH38* was increasingly induced in *btsl1x2* roots with a significant difference between the Zn-exposed half vs the non-exposed half of the root system (**Figure 5D**, 210- vs 80-fold, p > 0.01). The same phenomenon was seen for the *IRT1* transcript (**Supplementary Figure S10D**) and when half the *btsl1x2* roots were exposed to Fe deficiency (**Supplementary Figure S10C, D**). Thus, metal-induced derepression of Fe deficiency-regulated genes is stronger in roots that lack BTSL1 and BTSL2, indicating that BTSL proteins act locally.

Recently, IMA1 has been implicated in the shoot-to-root Fe signalling pathway (García *et al*., 2022; Grillet *et al*., 2018) and suggested to interact with BTSL proteins (Lichtblau *et al*., 2022). Consistent with this, and with its appearance in Cluster II of the RNA-seq analysis (**Figure 3C**), *IMA1* is induced to a similar level in both standard and Zn-exposed root halves in wild type (15- and 21-fold respectively) and in *btsl1x2* (7-fold and 4-fold, **Figure 5E**), further supporting **Figure 3** data that *IMA1* is upstream and unaffected by loss of BTSL function. Similarly, there was no significant difference in the level of *FER1* induction between standard and Zn-exposed roots in either genotype (**Supplementary Figure S10B**), as *FER1* is not BTSL-regulated. In summary, the split root data suggest that BTSL proteins act downstream of systemic Fe deficiency signals.

## DISCUSSION

Analysis of mutants that are tolerant to external Zn, such as the *btsl1x2* mutant, could help answer outstanding questions in Zn and Fe crosstalk. Specifically, how are Zn and Fe signals integrated and by which regulators and regulatory mechanisms? Previously suggested points of cross-talk and regulation, which are not mutually exclusive, are the Zn-induced internalization of IRT1; competition of Fe and Zn for protein binding sites; and disruption of systemic signalling mechanisms (Hanikenne *et al*., 2021). Our phenotypic, ionomic and transcriptomic analyses of the *btsl1x2* mutant in comparison to wild type suggest a model to explain why excess Zn causes systemic Fe deficiency (**Figure 6**) and how mutation of certain genes, in particular genes negatively affecting FIT activity (**Table 1**), suppress this phenomenon.

**Figure 6:**
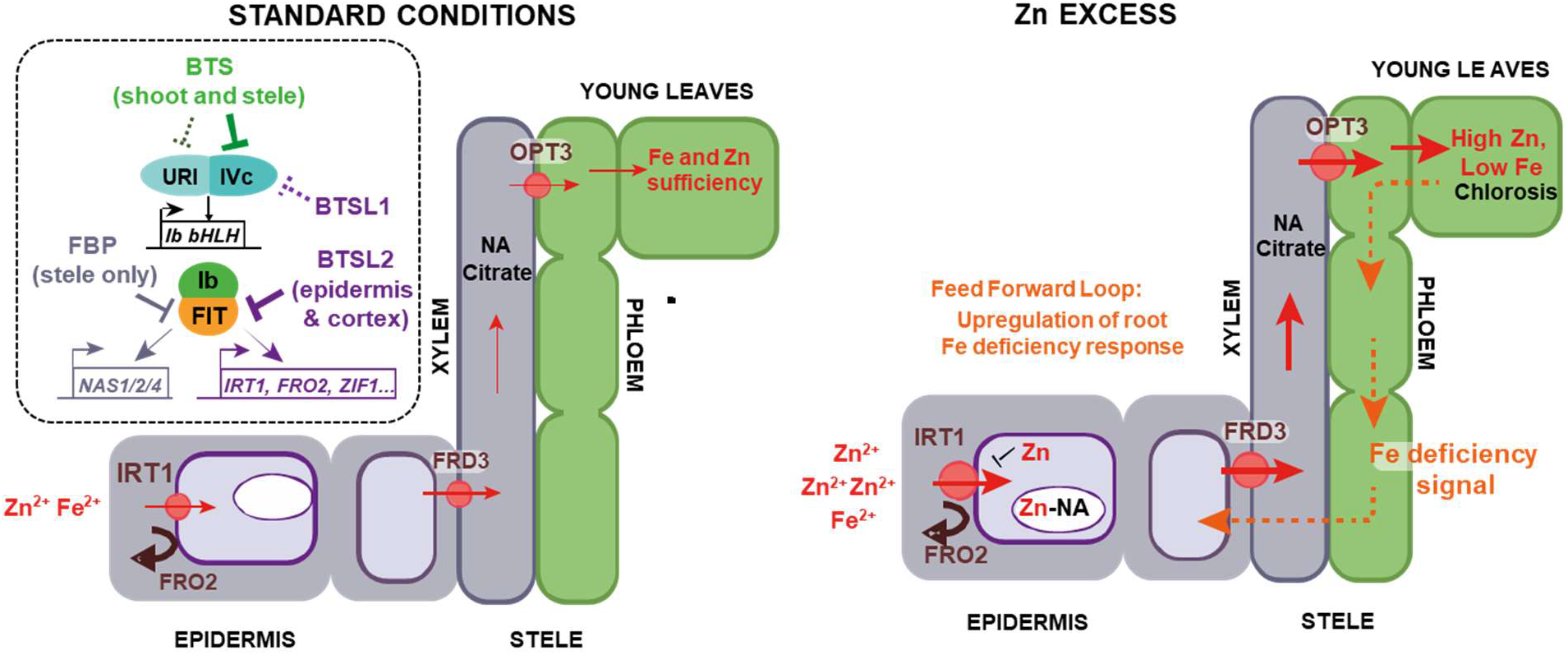
Model for *Arabidopsis* response to Zn excess. Under standard growth conditions (Fe:Zn ratios of approximately 5-25), BTS targets bHLH IVc transcription factors for degradation in the shoots and root stele (indicated by a solid line inhibition symbol), and possibly also bHLH121/URI (dashed line inhibition symbol). BTSL2 is thought to target FIT for degradation, and BTSL1 potentially targets IVc bHLH transcription factors, in the root epidermis and cortex. The effect of BTS and BTSL activity is to repress expression of Fe uptake and mobilisation genes, such as *IRT1, FRO2* and *ZIF1*, preventing accumulation of Fe, as well as Zn. FBP inhibits FIT-dependent induction of *NAS1/2/4* in the root stele. Under Zn excess conditions (Fe:Zn ratio of approximately 0.05), de-repression of bHLH121/URI, IVc bHLH and FIT leads to the upregulation of *IRT1* and *FRO2*, allowing increased uptake of Fe, but additionally of Zn as well. Increased Zn concentration in the cytoplasm leads to internalisation and thus removal of IRT1. Increased expression of *NAS* genes and *ZIF1* enhances vacuolar sequestration of Zn-NA in root vacuoles, whilst promoting Fe transport to shoots. Transport of Zn to the shoots results in a high Zn to Fe ratio, triggering chlorosis and an Fe deficiency signal from shoots to roots. The shoot-borne Fe deficiency signal creates a feed-forward loop, further upregulating metal uptake in wild-type plants.

In the presence of excess Zn, wild-type Arabidopsis seedlings take up large amounts of Zn and this is translocated to the shoots. Thus, IRT1 internalization cannot prevent the influx of Zn.

Increased expression of genes involved in vacuolar Zn sequestration (*MTP3, ZIF1* see **Figure 3C**) indicates that part of the internalized Zn is rendered non-toxic by storing it in vacuoles. However, impaired root growth (**Figure 2D**) suggests that not all excess Zn can be made harmless. Zn excess in the medium also leads to massive Fe accumulation in the roots (**Figure 2B**) but not to increased Fe in the shoots. Based on the transcriptomics data, it appears that more Fe is acquired by the roots as a consequence of the induction of the Fe deficiency response, but that the Fe cannot progress to the shoots. Our data is consistent with a possible deposition of Fe in the root apoplast, conceivable as a result of suboptimal FCR activity (**Figure 4**), or in other locations where Fe cannot be removed by EDTA washes. Because neither ferritin nor vacuolar Fe transporters are transcriptionally induced in response to excess Zn in wild-type roots (**Figure 3C**), this apoplastic deposition seems the most likely scenario, which could be verified by detailed microscopy studies of cellular Fe pools in the roots in the future.

Lack of BTSL1 and BTSL2 partially prevents Fe accumulation in Zn-exposed roots, which is counterintuitive given that Fe uptake genes are more highly induced than in wild type and no more Fe is translocated to the shoots. Possibly, slightly increased Fe uptake initially suppresses Zn-induced internalization of IRT1 (Barberon *et al*., 2014; Dubeaux *et al*., 2018; Martin-Barranco *et al*., 2020), allowing sufficient Fe uptake and subsequent transport to shoots. However, increased Fe uptake via IRT1 alone is not sufficient to confer Zn tolerance, and in fact the *idf1* mutant which is impaired in IRT1 turnover, shows Zn accumulation and sensitivity to Zn excess (Dubeaux *et al*., 2018). Interestingly, the *bts-1* mutant under Zn excess accumulates more Fe and Zn in both roots and shoots than wild type (Zhu *et al*., 2022). *BTS* is predominantly expressed in leaves, whereas *BTSL* expression is restricted to roots. *bts* mutants accumulate Fe in shoots (Hindt *et al*., 2017), whereas *btsl* mutants only accumulate Fe under specific conditions (Rodriguez-Celma *et al*., 2019). Thus, the difference in shoot Fe and Zn accumulation between *bts* and *btsl* mutants is likely associated to where the proteins are normally active.

In addition to the striking difference in root Fe accumulation between wild-type and the *btsl double* mutant, another key difference is the physiological Fe status of the shoots based on transcriptomics data (**Figure 3**). Wild-type shoots under Zn excess have only a slight decrease in Fe (**Figure 2B**) but are chlorotic (**Figure 1B)** and have high expression of genes normally induced by Fe deficiency (**Figure 3**; **Supplementary Figure S8**). By contrast, the *btsl1 btsl2* double mutant has near-normal chlorophyll levels and is Fe-replete according to transcriptomics analysis. This difference is not affected by exposing part of the root system to Zn excess (**Figure 5**).

Thus, it is possible that wild-type plants exposed to Zn excess become trapped in a feed-forward loop of impaired Fe uptake resulting in physiological Fe limitation in shoots and subsequent systemic Fe deficiency signalling to roots (**Figure 6**). This feed-forward cycle observed in wild type is similar to that seen in the *frd3* mutant, which shows constitutive activation of the root Fe deficiency response due to shoot-borne deficiency signalling, despite over-accumulating Fe in roots (Scheepers *et al*., 2020). This cycle can be partially broken at the level of FIT through mutations that lead to increased FIT protein levels, such as *btsl1 btsl2* (this study) and *fbp* (Chen *et al*., 2018), or mutations in upstream regulators such *as bts* (Zhu *et al*., 2022). It would be interesting to investigate whether natural variation in *BTSL* or *BTS* expression patterns, or ability to interact with transcription factor targets (FIT, IVc or IVb) or IMA proteins, impacts Fe and Zn phenotypes.

In addition, we propose that the tight regulation of FIT by BTSL1/2 is a trade-off between preventing Fe toxicity against preventing Zn toxicity under fluctuating nutrient availability. BTSL1/2 have a physiologically important role in rapidly removing FIT and switching off Fe uptake when roots are newly exposed to Fe after a period of deficiency. This mechanism prevents a flux of Fe into the plant that would be toxic. When BTSL1 and BTSL2 are absent, slightly enhanced Fe uptake is beneficial to outcompete with Zn for uptake and at protein binding sites, and the non-physiological Fe-deficiency response triggered by Zn excess is supressed. Thus, not having BTSL proteins, which are specific to dicotyledon plants, helps plants cope better with excess Zn in the environment, but it would be disadvantageous when Fe is in excess. It would be interesting to investigate natural variation on soils with contrasting Fe and Zn concentations, and see if there are accession that have lost functional BTSL proteins.

In summary, by regulating FIT and also FIT-independent transcriptional networks the BTSL proteins are part of an important control point in balancing Fe and Zn uptake and preventing toxicity.

## Abbreviations

bHLH: basic helix-loop-helix
BTS: BRUTUS
BTSL: BRUTUS-LIKE
FBP: FIT-Binding Protein
FCR: Ferric Chelate Reductase
Fe: iron
FIT: <u>F</u>ER-like Iron deficiency induced <u>T</u>ranscription factor
FRO2: Ferric Reduction Oxidase 2
FW: fresh weight
GSH: glutathione
HBED: N,N′-bis(2-hydroxyphenyl)ethylenediamine-N,N′-diacetic acid
HMA: Heavy Metal ATPase
IRT1: Iron-Regulated Transporter1
Mn: manganese
MTP: Metal Tolerance Protein
NA: nicotianamine
NAS: Nicotianamine Synthase
ZIF: Zinc-Induced Facilitator
Zn: zinc

## Supplementary data

***Table S1:*** qRT-PCR primers used in the study

***Dataset S1:*** RNA-seq data for the *btsl1x2* response to Zn excess

***Figure S1:*** The *btsl1 btsl2* phenotypic response to Zn excess

***Figure S2:*** The *btsl1 btsl2* phenotypic response to Mn deficiency and excess

***Figure S3*:** Principal component analysis (PCA) and comparison of the number of differentially expressed genes in the shoot transcriptomic response to Zn excess for wild type and *btsl1 btsl2*

***Figure S4*:** Gene Ontology (GO) analysis and comparison of number of differentially expressed genes in the root transcriptomic response to Zn excess in wild type and Zn excess ***Figure S5*:** Hierarchical clustering of differentially expressed genes in wild type and *btsl1 btsl2* roots in response to Zn excess

***Figure S6:*** Gene Ontology (GO) analysis of differentially expressed genes in wild type and *btsl1x2* shoots in response to Zn excess

***Figure S7:*** qRT-PCR verification of Fe deficiency-responsive genes in roots

***Figure S8:*** Hierarchical clustering of differentially expressed genes in wild type and *btsl1 btsl2* shoots in response to Zn excess

***Figure S9:*** Shoot phenotype and shoot expression of *bHLH39* from the split root experiment

***Figure S10:*** Root expression of *IMA1, FER1, bHLH38* and *IRT1* from the split root experiment

## Author contribution

CS, UK, DS and JB: conceptualization; CS and JR-C: methodology; CS: formal analysis; CS and MO: investigation; YL: data curation; CS and JB: writing - original draft; JR-C, UK and DS: writing - review & editing; CS and JB: visualization; JR-C and DS: supervision; UK, DS and JB: funding acquisition

## Data availability

The data supporting the findings of this study are available within the supplementary materials available online or from the corresponding author, Janneke Balk, upon request.

## Acknowledgements

We thank Graham Chilvers from the Science Analytical Facility, University of East Anglia, for ICP-OES analysis. We also thank Michele Oliva for preliminary studies supervised by Ute Kraemer (http://archiv.ub.uni-heidelberg.de/volltextserver/14879/1/thesis_Michele_Oliva.pdf)

